# Event-based vision sensor enables fast and dense single-molecule localization microscopy

**DOI:** 10.1101/2022.07.22.501162

**Authors:** Clément Cabriel, Christian G. Specht, Ignacio Izeddin

## Abstract

Single-molecule localization microscopy (SMLM) is often hampered by the fixed frame rate of the acquisition. Here, we present an alternative new approach to data acquisition and processing based on an affordable event-based sensor. This type of sensor reacts to light intensity changes rather than integrating photons during each frame exposure time. This makes it particularly suited to SMLM, where the ability to surpass the diffraction-limited resolution is provided by blinking events. Each pixel works independently and returns a signal only when an intensity change is detected. Since the output is a list containing only useful data rather than a series of frames, the temporal resolution is significantly better than typical scientific cameras. We demonstrate event-based SMLM super-resolution imaging on biological samples with spatial resolution on par with EMCCD or sCMOS performance. Furthermore, taking advantage of its unique properties, we use event-based SMLM to perform very dense single-molecule imaging, where frame-based cameras experience significant limitations.

---

The advent of single-molecule localization microscopy (SMLM) brought a vast improvement in image resolution [1, 2, 3] and has become an invaluable tool for cell biology as it can resolve nanometric cellular structures. Significant advances have been possible through the development of new optical techniques [4, 5], labeling methods [6, 7, 8] and the synthesis of brighter dyes compatible with live-cell imaging [9, 10]. For recent reviews of the field of SMLM, refer to [11, 12]

Retrieving the positions of single molecules is pivotal in applications as varied as 3D imaging [13, 14, 15, 16], spatial analysis of protein clusters [17, 18] or protein dynamics in the cell [19]. Still, the point-spread function (PSF) carries significant additional information, giving access to the spectrum of the dyes for multicolor imaging [20, 21], the emitter’s orientation [22], the local environment of the molecules through modifications of the fluorescence intensity [23, 24], and fluorescent state lifetime [25, 26, 27, 28].

Regardless of their use, single-molecule experiments are routinely limited by the acquisition speed. On the one hand, SMLM experiments often require more than 20000 frames, i.e. 10 minutes with 30 ms exposure time. On the other hand, the chosen exposure time sets a hard limit on the temporal scale at which fast dynamic processes are observable. While the importance of temporal resolution is obvious in live cells, it should not be overlooked in experiments with fixed samples. More generally, increasing the acquisition rate allows more efficient data collection, which can be important to better describe biological phenomena and improve statistical significance. In some cases, it can even be used to develop automated high throughput data collection setups and analysis workflows [29, 30].

Improving the data recording rate is no small task, however, as frame-based scientific cameras (EMCCD and sCMOS) work at a fixed frame rate chosen before the acquisition. This raises serious issues whenever the spatial distribution of proteins in the sample is heterogeneous, or when different targets in the experiment obey dynamic processes at different time scales. In any case, the frame rate typically has to be adapted to the fastest processes or the highest densities, which leads to a detrimental increase of noise at best or a loss of relevant information at worst.

On the other hand, event-based sensors (also called neuromorphic sensors) use a very different principle. An event-based sensor is an array of pixels functioning independently, and sensitive to intensity changes rather than to an average irradiance over a fixed exposure time. Although each pixel has a dead time in the range of a few μs to 100 μs, it does not work at a fixed frame rate. Instead, it returns an ‘event’ after it detects a change of irradiance. Therefore, the sensor essentially returns only meaningful data stored as a list of events, which potentially allows a vast increase of data collection rate compared to scientific cameras. As a comparison, the fraction of camera-recorded data effectively used for localization purposes varies from around 10 % in the case of dense dSTORM experiments to less than 0.1 % in sparse SPT acquisitions. Event-based sensors generally have pixel numbers of the order of a million and a pixel size of the order of 10 μm, and are more affordable (<6000 $) than sCMOS and EMCCD cameras.

Here, we propose an implementation of SMLM experiments with an event-based sensor. We describe the principle of event-based sensing and the data processing steps. We characterize the response of the sensor to a fluorescent signal and assess the localization precision. We furthermore demonstrate the detection of organic dyes for single-molecule fluorescence imaging and use it to produce a super-resolution image of a fixed immunolabeled sample. Finally, we use the unique features of event based sensing to improve localization performance in the high density regime where PSF overlap cause classical frame-based SMLM to fail.

## Results

### Principle and implementation

For general reviews about event-based sensing, readers may refer to [31, 32]. As all the pixels of the array are independent, the response of an event-based sensor can be described through the response of one pixel, as shown in **Fig. 1a**. Each pixel returns a signal when variations of intensity, positive or negative, exceed a given threshold or sensitivity value *B*, common to all pixels. *B* is set by the user to define the threshold as illustrated in **Fig. 1b**. The intensity variation detection is logarithmic according to the manufacturer specifications, and the sensitivity can be as low as 11 % [32]. Whenever an event is triggered, the sensor returns a signal containing the *x* and *y* pixel coordinates, the polarity (positive or negative) and the timestamp, and the new intensity value is set as the new reference level. A large variation of the signal causes the pixel to return several events in a quick succession, which readily provides a certain *quantitativity*. A more detailed description of the response of each pixel is provided in **Supplementary note 1.**

**Figure 1:**
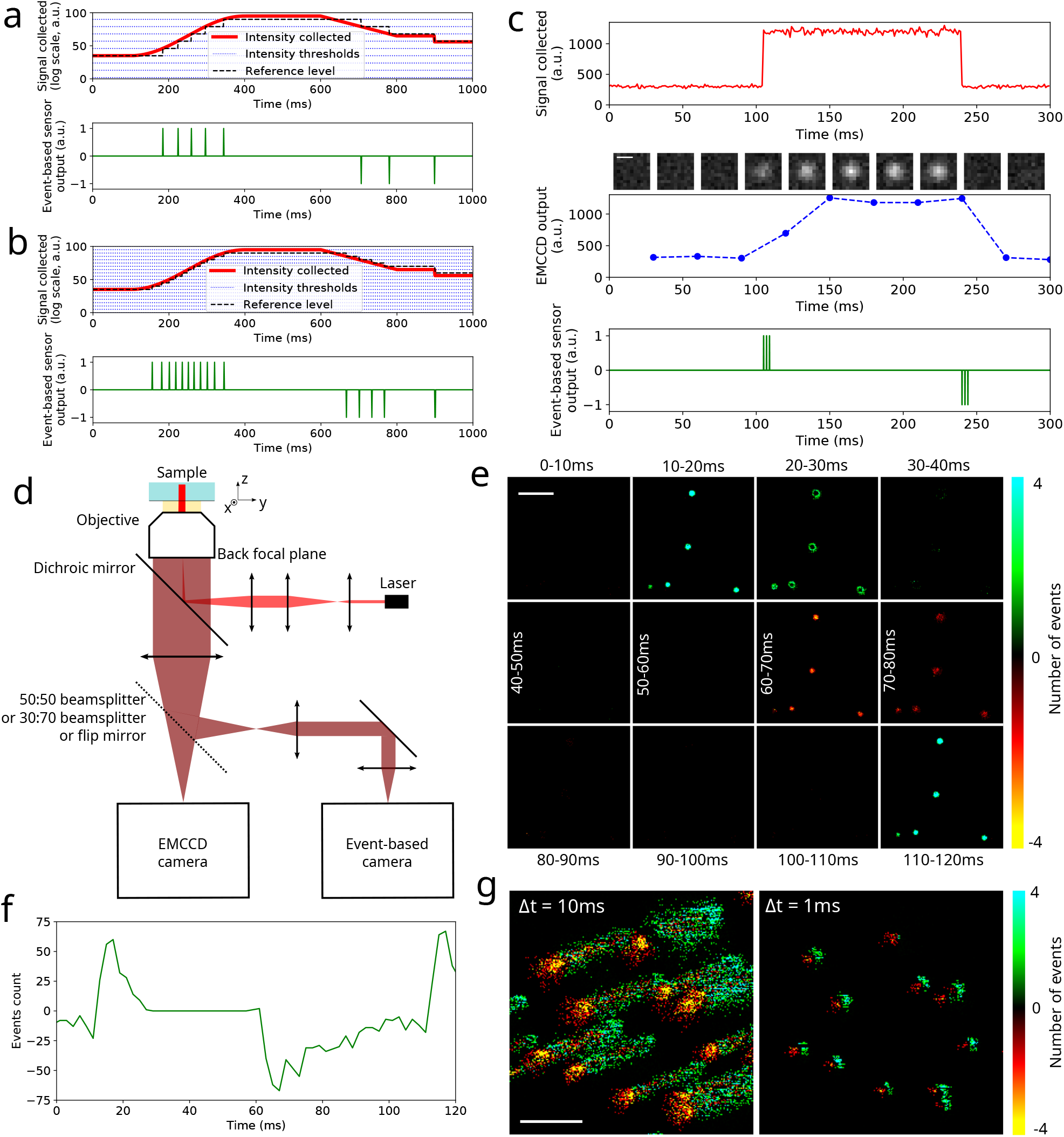
Principle of the event-based detection. **a** Description of the working principle of a pixel in an event-based sensor. A given optical intensity signal *I*(*t*) is assumed, the sensitivity value *B* is set to an arbitrary value. The reference value is also displayed, as well as the response from the pixel. **b** Influence of the sensitivity on the output signal for the same *I*(*t*) as in **a**, but *B* is lower (i.e. the detection is more sensitive). Therefore, the number of events generated is higher. **c** Comparison of the responses produced by a scientific camera and an event-based sensor. The top row shows a theoretical *I*(*t*) profile emitted by a fluorescent molecule turning on and then off, the middle row is the signal acquired by an ideal camera working at a 30 ms exposure time (the corresponding frames are displayed above the graph), and the bottom row displays the signal returned by an event-based sensor with arbitrary sensitivity. Optical setup used for the single-molecule experiments. A laser produces a Gaussian excitation beam. The detection path comprises two detectors: and EMCCD camera and an event-based sensor. We use switchable deflectors to alternate between the two detection paths: either a 50:50 beamsplitter, a 30(EMCCD):70(event-based) beamsplitter, a mirror or nothing. **e** Frames generated from the events acquired in an experiment where 200 nm fluorescent beads on a coverslip are excited with a square modulated excitation with a period of 100 ms. The frames are generated with a time bin of Δ*t* = 10 ms. The color codes the number of events in each pixel, with respect to the polarity (positive or negative). The time limits are written for each frame. **f** Event time profile extracted from a single bead from **(e)** with a Δ*t* = 2 ms time bin. The number of events represented corresponds to the sum of all the pixels over the PSF. **g** Frames generated at different time bins from an acquisition of 40 nm fluorescent beads deposited on a coverslip while the stage is translated from the bottom left to the top right to simulate molecule motion. From the same event list, frames are generated at Δ*t* = 10 ms (left) and Δ*t* = 1 ms (right). Scale bars: 0.5 μm (**c**), 5 μm (**e**), 5 μm (**g**).

**Fig. 1c** illustrates the differences between event-based and classical cameras. First, the event-based sensor detects only intensity changes (in our case, a molecule turning on or off, or the motion of a moving emitter). This offers very promising perspectives when it comes to implementing live data processing and image reconstruction, or high-throughput automated experiments. Another difference is the dynamic range, which is vastly superior for the event-based sensor, making it almost impossible to saturate the sensor with fluorescence signals, which can be beneficial when using fiducial markers. Finally, the event-based sensor does not use a fixed frame rate and thus does not return a stack of frames. This implies that the temporal resolution of the acquisition is not limited by user-defined parameters. Since the temporal bandwidth is not infinite (each pixel has a certain dead time in the range of a few μs to a few hundreds of μs [33]), the time resolution is determined in the processing stage by the time base used for the data treatment. This feature is very interesting for SMLM imaging as it allows to process the same dataset with different time bases to investigate biological processes at different time scales.

We used such an event-based sensor in an optical setup designed for SMLM experiments presented in **Fig. 1d**. The detection path is built to offer some modularity by distributing the fluorescence light in either one or both of the sensors, the EMCCD and the event-based sensor (see **Methods**). We propose two complementary data display approaches. A purely frame-based approach is shown in **Fig. 1e**. Frames are generated by binning all the events in a certain time interval Δ*t* in a pixelized canvas (each positive event is counted as +1 and each negative as –1). This visualization method emphasizes the spatial distribution of events at the expense of some time resolution. We represent data acquired on fluorescent beads on a coverslip with 10-Hz square modulated excitation with a chosen Δ*t* = 10 ms frames. The rising and falling edges of *I*(*t*) (frames 10–20 ms and 60–70 ms respectively) are clearly visible, as well as the near absence of output signal when the intensity is constant, whether the source is emitting (30–60 ms) or not (80–110 ms). Finally, one can note on frames 20–30 ms and 70–80 ms that the signal is not returned instantly—more specifically, the dimmer pixels at the edge of the PSF respond more slowly, which explains the visible rings (we attribute this to the time necessary to sample the photons into an intensity measurement). **Fig. 1e** furthermore highlights the quantitative aspect of the detection—the bead at the bottom left is less bright and thus generates fewer events.

In **Fig. 1f**, we propose a complementary data display based on temporal profiles over all the pixels of a PSF, which highlights the nature of the signal returned by the event-based sensor. Although this representation also relies on time binning, it is typically done with much lower values of Δ*t*, which is more suited for the study of the time response of single molecules. This comes at the cost of a certain loss of spatial information, as all the pixels in a PSF are summed together. **Fig. 1f** incidentally reveals that negative events seem on average slower than positive events. We do not have an explanation for this effect, but we assume it can be addressed by tuning the electronics coupled to the sensor.

Although the temporal bandwidth is not defined by the acquisition parameters, the processing algorithms have to be carefully designed to account for the time scales that match the desired biological application. We highlight this in **Fig. 1g** by moving fluorescent beads with the piezoelectric stage. A small time bin (Δ*t* = 1 ms) efficiently samples the dynamic to reveal two opposed lobes (leading and trailing edges) at the cost of a relatively modest number of events. On the contrary, increasing the time bin to Δ*t* = 10 ms provides a large number of events for a better statistical set, which is spoiled by a dramatic motion blur as well as a loss of resolution. Obviously, the value of the time bin has to be matched to the phenomenon under investigation, but no knowledge or assumption is required prior to the acquisition (contrary to scientific camera-based approaches), as the selection of Δ*t* is done as part of the processing workflow. This could even be used to perform a multi-timescale processing on the same dataset to extract both slow and fast processes.

### Single molecule localization

We then set out to detect and localize single molecules, focusing on blinking labels as in (d)STORM, PALM or DNA-PAINT microscopy. Our processing method for single-molecule localization is described in **Supplementary note 2.** As a first step, we assessed the localization precision with fluorescent beads on a coverslip. The principle of this measurement is summarized in **Fig. 2a** as well as in the **Methods** section, and the precision results are displayed in **Fig. 2b**, along with the Cramér-Rao Lower Bound (CRLB) values for the experimental parameters obtained with the EM-CCD camera. Overall, the event-based sensor precision was found to be on par with the EMCCD center of mass calculation, or slightly better. In particular, precisions below 10 nm are found in the typical levels of signals of SMLM fluorophores, and sub-5 nm precision is obtained with approximately 5000 photons, which is the level of signal emitted by many far red organic dyes such as Alexa Fluor (AF) 647. Precision tends to improve slightly with finer sensitivity values. Interestingly, experimental precision was found to be slightly better than the EMCCD CRLB, which we attribute to very low levels of background-induced events and read noise when working with an event-based sensor.

**Figure 2:**
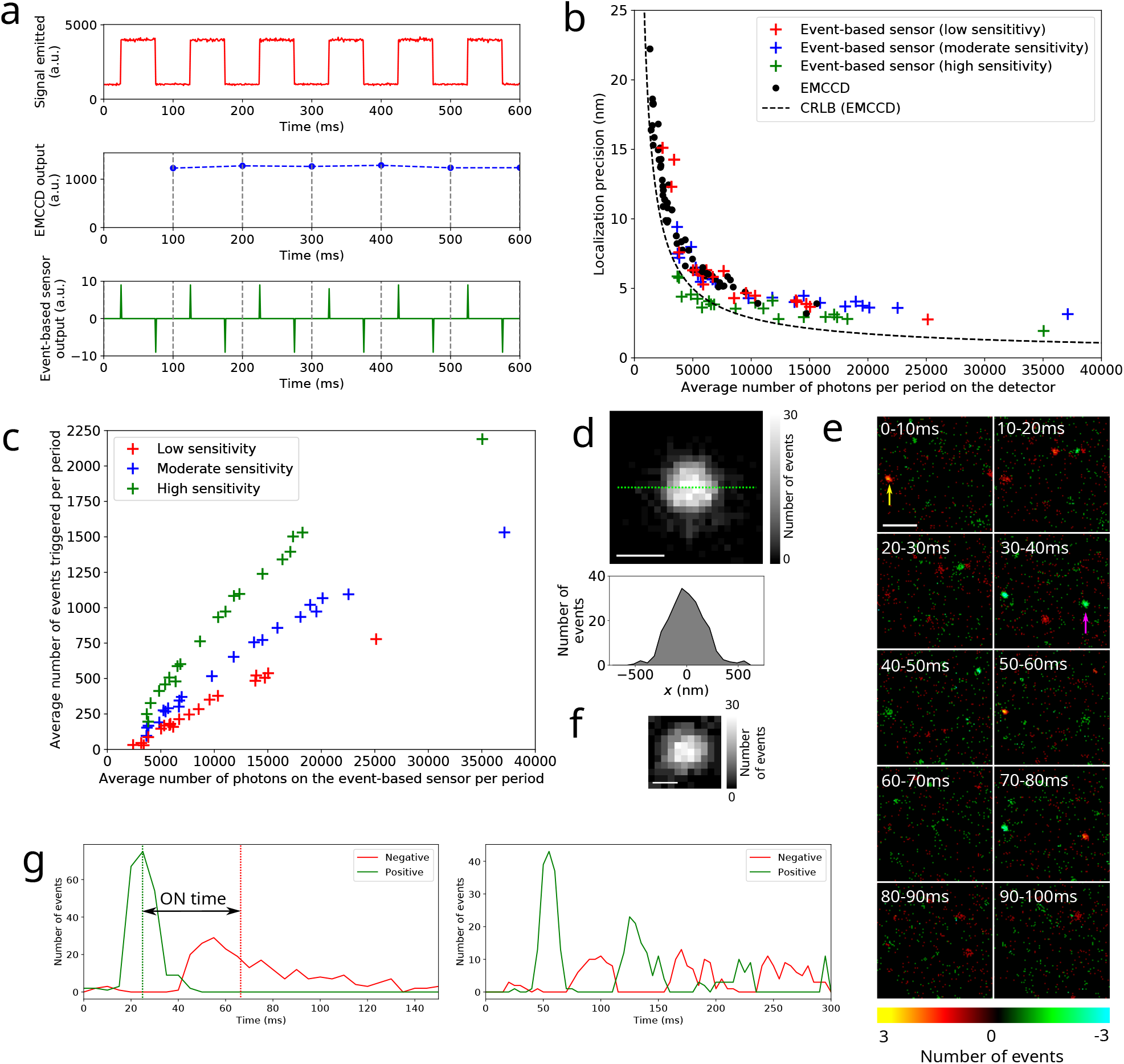
Single molecule localization and characterization of the performance. **a** Schematic of the experiment used to measure the localization precision. Fluorescent beads deposited on a coverslip are illuminated with a square modulated laser at 10 Hz. The EMCCD exposure time is set to 100 ms, i.e. exactly one period, so the signal is constant over all the frames (with the exception of the photon noise). The event-based sensor detects the rising and falling edges, and all the events in one period are pooled to compute the center of mass position. **b** Localization precision results displayed as a function of the number of photons incoming on each of the detectors (EMCCD or event-based sensor) per period (calculated from the number of photons detected on the EMCCD taking into account the 30:70 ratio). The results for the event-based sensor are displayed for different levels of sensitivity. The CRLB curve corresponding to the EMCCD detection is also shown. **c** Linearity of the response, shown for different levels of sensitivity. The total number of events detected per period is displayed as a function of the number of photons incoming on the event-based sensor per period. Note the quasi-linearity of the response in this signal range. **d** Top: PSF reconstructed by summing the events (regardless of their sign) detected over one period of 100 ms for one bead. Bottom: *x* profile along the green dashed line. **e** Frames reconstructed with Δ*t* = 10 ms from a sample of AF647 deposited on a glass coverslip excited with a continuous laser in a dSTORM buffer to induce blinking. Note that this data display reveals very different blinking behaviors from one molecule to the other—the molecule indicated with a yellow arrow undergoes multiple blinking events in a quick succession while that indicated by a magenta arrow exhibits a much simpler behavior with essentially one rising edge and one falling edge. **f** PSF reconstructed from all the events (both positive and negative) in one AF647 blinking event. **g** Time profiles plotted for two different AF647 molecules taken from the same acquisition as e (but not those shown with the arrows). The one on the left shows a simple blinking profile, which allows the calculation of the ON time, defined as the difference of the mean time of the negative events and the the mean time of the positive events. The profile on the right, on the contrary, displays a complex behavior with multiple blinking events. Scale bars: 500 nm (**d**), 2 μm (**e**), 250 nm (**f**).

We furthermore characterized the linearity of the response, which is important for the quantitative aspect of the output data. We displayed the number of output events per period as a function of the number of incoming photons per period in **Fig. 2c**. As expected, the number of events recorded for a given input signal increases as the sensitivity gets finer (i.e. from low to high sensitivity). Interestingly, the relationship between the output and the input was found to be largely linear, unlike the specifications given by the manufacturer. We hypothesize that the output becomes logarithmic at higher levels of input signal (exceeding the typical range of SMLM intensities), as this particular sensor has been originally designed to work at ambient light levels.

To further illustrate the capability of the sensor for SMLM, we reconstructed and displayed the PSF obtained over one period in **Fig. 2d** for an input signal of 10000 photons and an integrated output signal of 1000 events approximately (with a high sensitivity). The PSF has a very regular shape, and a profile plot yields a width *w* around 190 nm (standard deviation), slightly above EMCCD value (*w*=170 nm, data not displayed).

After the characterization of the system with fluorescent beads, we set out to use the event-based sensor to detect single molecules. We performed acquisitions with AF647 dyes on a coverslip (see **Methods**) with the event-based sensor, and reconstructed frames at Δ*t*=10 ms (**Fig. 2e**). Their blinking can be efficiently monitored—molecules exhibit a typical positive rising edge, followed by a few tens of ms without any signal, and finally a negative falling edge at the same position as the rising edge. Again, the PSF displays a very similar aspect to camera-based PSFs (**Fig. 2f**).

We also investigated the temporal behavior of the fluorophore blinking, which is readily returned by the sensor output (**Fig. 2g**). While some dyes have a very simple profile with well-defined rising and falling edges (the falling edge still displaying the tail mentioned previously) and few events in between, others display a very erratic behavior, with a succession of multiple rising and falling edges, and seemingly intermediate levels of gray. This is consistent with the blinking regime observed on an sCMOS camera under the same illumination conditions and with an exposure time of 10 ms (data not displayed). Organic dyes in oxygen reduction buffers are known to have a complex photophysical behavior, often far from a clean square signal [34]. We point out that, interestingly, event-based SMLM could provide an efficient way to both characterize these temporal emission fluctuations and image the dyes without any loss in performance or information.

### Event-based SMLM bioimaging

As organic dyes could be efficiently detected with our event-based single-molecule localization method, we moved on to imaging fixed biological samples labeled with AF647 targeting the tubulin cytoskeleton in fixed COS-7 cells (see **Methods**). Using a 50:50 beamsplitter in the detection path, we performed a simultaneous EMCCD and event-based acquisitions. The processing is described in **Supplementary note 2.**

2D SMLM images obtained with the event-based and the EMCCD sensors were found to be very similar at the scale of the whole field of view (**Fig. 3a**). At the nanometer scale (**Fig. 3b**), the event-based detection performed on par or even outperformed the EMCCD, allowing to resolve the hollow core of the microtubules (see the cyan arrows and the corresponding profiles in **Fig. 3c**). Note that taking into account the diameter of the microtubules (30 nm) and the size of the antibody labeling (10 nm on each side), the apparent diameter is expected to be around 50 nm [8, 15]. While both the event-based center of mass calculation, the event-based Gaussian fitting and the EMCCD Gaussian fitting reveal the apparent hollowness of the microtubules, a simple center of mass calculation with the EMCCD fails to do so. We furthermore performed resolution assessments using Fourier Ring Correlation (FRC) [35, 36]. **Supplementary Fig. 1** shows an event-based resolution of 36 nm with a center of mass calculation and 30 nm with Gaussian fitting, very close to the reliable Gaussian fitting EMCCD resolution (28 nm), and significantly better than a simple center of mass calculation on the EMCCD data (50 nm). All the resolution measurements are summarized in **Supplementary Table. 1.** While using a dual-view detection setup is useful to monitor the acquisition in real time and to compare the final results, it is not necessary and it reduces the number of collected photons. To exploit the full potential of the event-based sensor, we performed an acquisition using a mirror in place of the beamsplitter cube (**Fig. 3d**). Again, the event-based SMLM ensures consistent localization performance over the whole field of view (with a resolution of 24 nm), and yields excellent precision that allows to resolve the microtubules’ apparent hollow core even more clearly (**Fig. 3e**).

**Figure 3:**
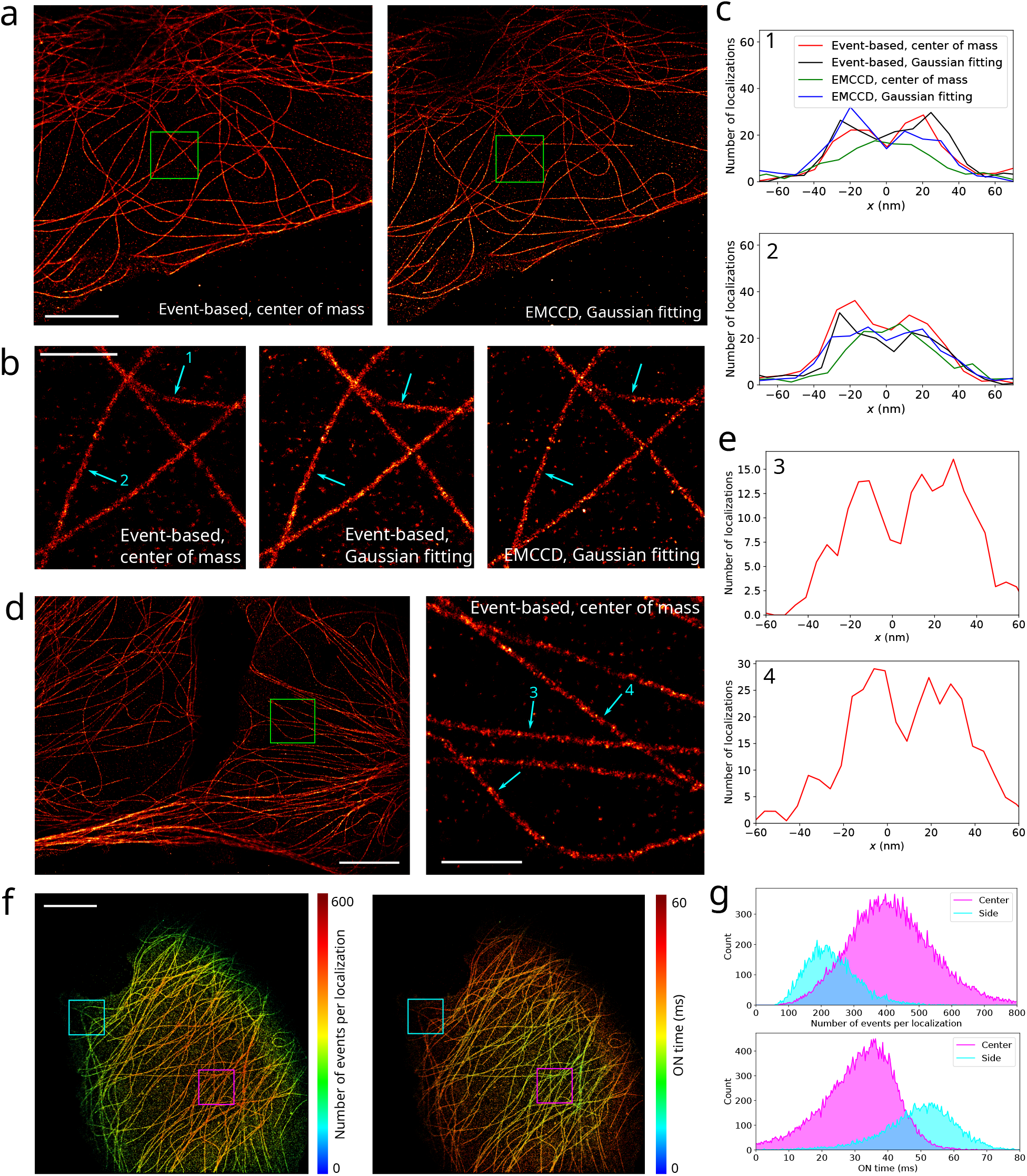
Imaging of fixed COS-7 cells labeled with AF647 against *α*-tubulin (see Methods). **a** Simultaneous 2D SMLM image (the density of molecules is color-coded) obtained with the event-based sensor and the EMCCD camera using a 50:50 beamsplitter in the detection path. **b** Zoom on the region of **a** indicated with a green square. We compare the resolution obtained with the event-based sensor (localization performed by center of mass calculation or Gaussian fitting) and with the EM-CCD camera (localization performed by Gaussian fitting; center of mass was calculated but is not displayed). **c** Molecule density profiles plotted perpendicular to the microtubules 1 and 2 in **b**. **d** 2D event-based SMLM image obtained with 100 % of the photons incoming on the event-based sensor. A zoom on the region indicated with a green square is also shown. **e** Molecule density profiles plotted perpendicular to the microtubules 3 and 4 in **d**. **f** Maps of the number of events per localization and of the ON time (color-coded values) for a different acquisition. **g** Corresponding histograms displayed for the two regions indicated with cyan and magenta squares. Scale bars: 5 μm (**a**, **d** left, **f**), 1 μm (**b**, **d** right).

The above results demonstrate the feasibility of SMLM bioimaging with an event-based sensor, which is on par with a conventional and more costly EMCCD camera. Moreover, our event-based method allows us to extract additional information other than the positions of detected molecules, such as photophysical properties of the fluorophores. **Fig. 3f** displays super-resolved maps of the ON time (as defined in **Fig. 2g**) and number of events detected for each molecule (which is closely related to the intensity of the molecule), and a more quantitative assessment of this is provided in **Fig. 3g**. The Gaussian shape of the illumination profile is clearly visible, and, as expected, both follow an inverse evolution. These quantities are often used to highlight chemical affinities or near-field optical effects at the molecular scale. For example, engineered DNA-PAINT strands can be used for multi-species demixing (using the binding time and frequency, as well as the intensities collected in three different color channels) [37]. Examples of the use of the ON time and intensities in single-molecule nanophotonics include probing the changes in photophysical properties that fluorescent molecules undergo near a gold surface or in a nanometric volume where the excitation field is vastly enhanced [38, 23].

### Event-based high density imaging

In order to exploit the unique capabilities of event-based sensing for single-molecule imaging, we use it in a regime where the density of PSFs is very high and their overlap significantly compromises the localization performance of frame-based acquisitions. The most common approach in classical SMLM with frame-based cameras is to adapt the density of active fluorophores to the most dense structures in the field of view to achieve sparsity, at the cost of potentially long imaging times. However, in some cases such as acquisitions in living samples, this may not be possible, resulting in under sampled structures. Very few techniques are available to tackle such situations. While SOFI [39, 40] yields consistent performance over a large range of molecule densities, it is not intrinsically a single-molecule approach, therefore it is not very well suited for quantitative applications such as molecule counting measurements. Other single-molecule approaches to process dense PSFs may require cumbersome procedures of deconvolution [41] or deep learning agorithms [42].

We propose to take advantage of the temporal resampling capabilities of event-based sensors to perform high density imaging. Indeed, with a standard camera, each molecule prevents the localization of other molecules in a diffraction-limited vicinity as long as it does not turn off. If that ON time is high with regards to the required density, the single molecule regime may be impossible to achieve independly of the framerate. In the same situation, an event-based sensor captures only the moments when that molecule turns on or off, allowing other blinking events in the same area during the ON cycle without resorting to complex algorithms.

To illustrate this principle, we performed acquisitions where a very dense layer of 488-nm fluorescent beads is deposited on a coverslip and excited with a constant power, while a few, sparse 647-nm beads are excited with a square-modulated 638-nm laser at 10 Hz. The results presented in **Fig. 4a** yield EMCCD frames where most 647-nm beads cannot be localized with usual approaches. Note in particular that beads 4 and 5 are very challenging to localize (or even visualize) among the other PSFs with standard processing algorithms. In contrast, the event-based reconstructed frames contain almost no trace of the static background (overlapping PSFs), and reveal very clearly the modulated beads.

**Figure 4:**
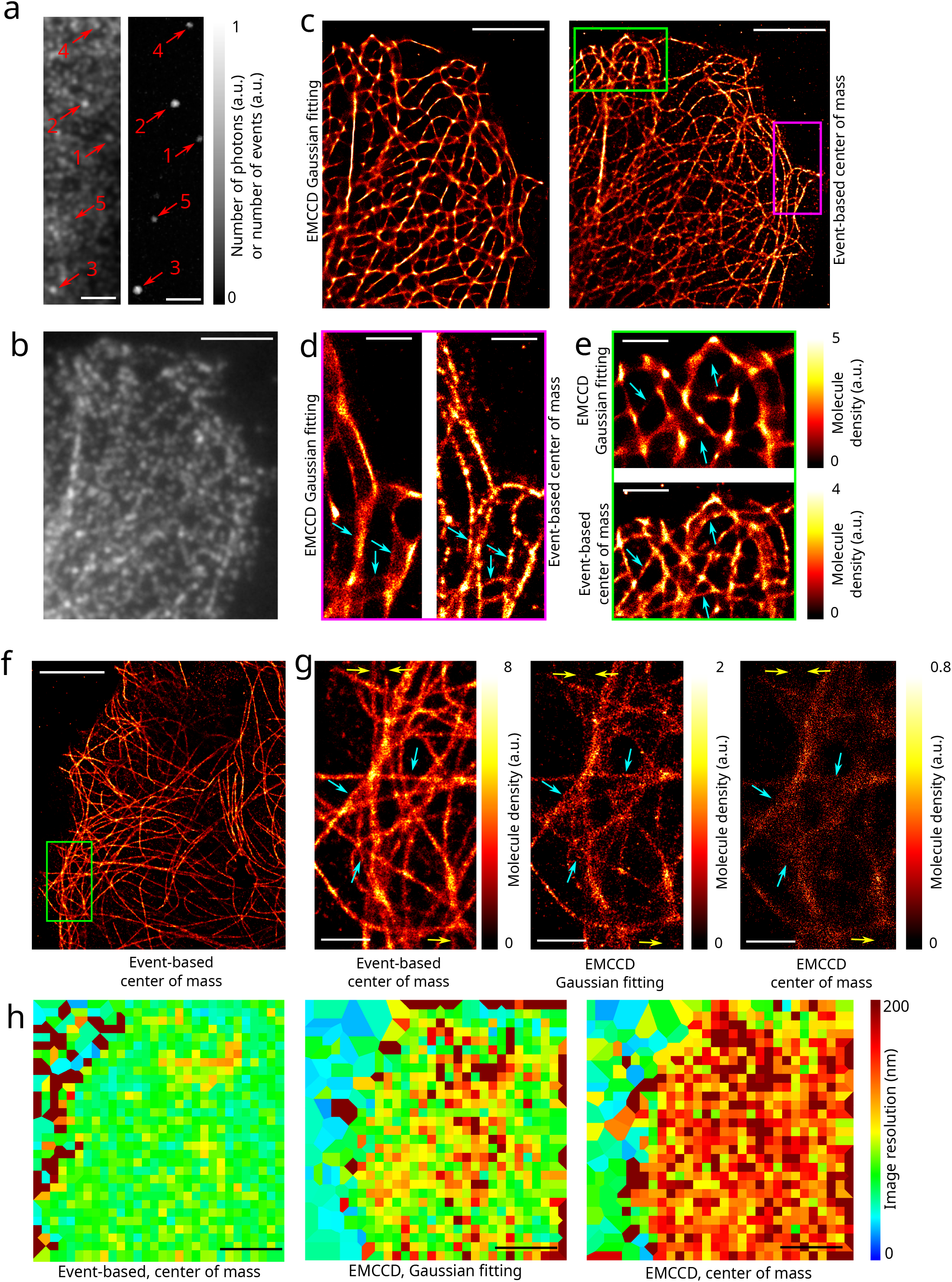
High density bioimaging. **a** Images of dense 488 nm and sparse 647 nm fluorescent beads illuminated with continuous 488 nm excitation excited and square modulated 638 nm excitation, using a multiband beamsplitter to collect both wavelength channels, and with a 50:50 beamsplitter between the two sensors. Left: Integrated signal (i.e. sum of all the frames) obtained with the EMCCD camera; right: integrated signal (i.e. sum of all the events) obtained with the event-based sensor. Some of the 647 nm beads are indicated with red arrows. **b** Raw sCMOS frame (50 ms exposure) showing the extreme density of molecules used in the high density DNA-PAINT experiments (fixed COS-7 cells, DNA-PAINT Atto655 labeling against *α*-tubulin). **c** Event-based and sCMOS SMLM images (the density of localizations is color-coded) obtained from the same high density DNA-PAINT experiment. **d–e** Zooms on two different regions of c indicated with magenta and green rectangles respectively. Cyan arrows highlight the better efficiency of event-based detection in terms of resolution in **d** and of fidelity of the reconstruction in **e**. **f** Event-based dSTORM image obtained from a 250 s long acquisition (fixed COS-7 cells, AF647 labeling against *α*-tubulin). **g** Zoom on the area indicated with a green rectangle in f, for the same 250 s long acquisition with simultaneous event-based sensor and EMCCD detections. The SMLM images are shown for different localization conditions: from the event-based sensor data with center of mass calculation (left), from the EMCCD data with Gaussian fitting (center), and center of mass calculation (right). Note the different scales of the colorbars. Cyan arrows indicate regions where the resolution is significantly better in event-based SMLM, and yellow arrows highlight missing structures on the EMCCD images that are well captured in event-based SMLM. **h** FRC resolution maps calculated on the area displayed in f for the different localization methods tested. Scale bars: 2 μm (**a**), 5 μm (**b, c, f, h**), 1 μm (**d, e, g**).

We then moved on to real blinking acquisitions in biological samples. Using the same setup as previously, we imaged fixed cells with very high DNA-PAINT imager concentration (see **Methods**). A raw 50-ms frame extracted from the EMCCD blinking movie presented in **Fig. 4b** highlights that the acquisition fails to achieve the single molecule regime. In such a situation, classical frame-based algorithms perform poorly, which results in a large degradation of the resolution. On the contrary, the event-based sensor is particularly helpful to temporally resample the signal, effectively decreasing the density of PSFs, as explained in more detail in **Supplementary note 3.** The adaptations made to the localization code for high density bioimaging are discussed in **Supplementary note 2.** As a consequence, the event-based localization image displayed very convincing resolution, and a much richer information content. This is visible in **Fig. 4c** as well as in **Fig. 4d**, where the arrows show well resolved pairs of microtubules that the EMCCD fails to resolve. Aside from the mere question of image resolution, high density acquisitions tend to induce significant reconstruction artifacts, a problem that is convincingly illustrated in [36]. **Fig. 4e** further stresses the better consistency of event-based SMLM compared to the EMCCD acquisition, as some microtubules are missing in the frame-based reconstruction (indicated by the arrows), which decreases the image information content.

As DNA-PAINT acquisitions are usually done on fixed samples with a controllable imager density, they rarely need high emitter densities. This does not apply to PALM and (d)STORM, however. The former is routinely used for living biological samples, where the acquisition speed is crucial and increasing the density of simultaneously active PSFs is tempting to increase the spatio-temporal resolution. While the latter can also be used to image living samples in certain cases [43, 44, 45], it has furthermore been shown that nanometer-range interactions between organic dyes lead to a vast number of blinking cycles in the pumping phase of dSTORM experiments (i.e. when the laser is set to high power at the beginning of the acquisition to switch molecules to the OFF state and achieve single molecule regime) [34], therefore wasting a large part of their initial photon budget during a time interval that is often discarded due to the very high density of emitters. For these reasons, we recorded the dSTORM pumping phase with both the event-based sensor and the EMCCD camera, using the the same AF647-labelled fixed sample as in **Fig. 3.** We chose a region where the microtubules are closely interwoven to ensure a high PSF density, and the acquisition was started at the same time as the laser excitation. Using a 50:50 beamsplitter cube, we compared EMCCD and event-based performances over the course of a short 250 seconds acquisition (during which the density of PSFs remains high).

Processing that short event-based dataset produced a strikingly well-sampled 2D SMLM image (**Fig. 4f**). The image exhibits a very uniform resolution, with little loss of sampling density or resolution either in dense areas or at the edges of the field of view. A more detailed view (**Fig. 4g**) reveals high resolution, good structure sampling and image fidelity, especially given the suboptimal acquisition conditions—50 % of the photons only, 4 minutes of acquisition compared to usual times around 20-30 minutes in dSTORM experiments, and high density of PSFs. Gaussian fitting event-based SMLM (super-resolved image not displayed) yielded very similar results as center of mass calculation. The image quality is even more striking compared with the EMCCD acquisition processed with a standard algorithm. **Fig. 4g** shows an undersampled SMLM image with a Gaussian fitting EMCCD-based processing that does not allow to properly distinguish a certain number of tubulin filaments (cyan arrows), especially in dense areas, as well as missing filaments (yellow arrows). A more basic EMCCD center of mass algorithm gave even worse results, with resolutions hardly better than the diffraction limit in dense areas. A quantitative assessment of the image resolution (**Supplementary Table 1)** yielded 60 nm for event-based SMLM (center of mass and Gaussian fitting), compared with 92 nm and 136 nm for EMCCD-based SMLM with Gaussian fitting and center of mass calculation, respectively. The resolution maps presented in **Fig. 4h** highlight not only the better overall performance of event-based SMLM, but also the consistency of the resolution, which is much more uniform and less sensitive to the density of the imaged structure compared with EMCCD acquisitions.

The high density localization performance could be further enhanced by refining the (currently basic) processing pipeline. In particular, adapting the processing conditions to the local density of molecules (in space and in time) could improve both the processing time and the localization performance. This is conceptually reminiscent of recent works such as [46], where the size of the spatial range available for PSF shaping is adapted in real time according to the density of molecules (which varies in time during the acquisition), or [47, 48], where the acquisition speed and illumination power are adapted in real time so that it allows recording of data at a sufficient rate while minimizing photo-bleaching, as well as saving storage and processing power. Nevertheless, a fundamental advantage of event-based SMLM is that the sensor records data as fast as it can, which means that no prior knowledge about the acquisition behavior is needed, and that workflow refinements are inherently processing-based. Another advantage is that, while the exposure time of cameras can be varied in time, all the chip is constrained with the same exposure time—by contrast, all event-based processing parameters can be varied along spatial dimensions as well.

## Discussion

We have demonstrated the use of event-based sensors for SMLM super-resolution fluorescence imaging. Besides their more affordable prices than conventional scientific cameras, their unique characteristics make them perfectly suited for detecting sparse events in space and time—incidentally the working principle of SMLM. With our event-based SMLM approach, we have obtained dSTORM super-resolution images on par with the widely used approach using a scientific camera. We further-more extracted quantitative photophysical information like the ON time and the number of events of the detected single fluorophores. Importantly, event-based SMLM opens new avenues for single-molecule applications where classical frame-based cameras are unable to perform, as we have shown in the case of high-density acquisitions with overlapping PSFs.

The range of applications of event-based sensing is broad since it is very well suited to the extraction of the dynamics of biological processes. In the context of imaging, it could be especially useful to monitor processes that exhibit a wide range of dynamic scales. An extreme example would be that of a system that would not evolve for several hours before suddenly exhibiting dynamics in the millisecond range. One can cite colloidal glass transition, where the dynamic ranges over more than ten orders of magnitude, with large variations over the field of view [49, 50].

In the framework of SMLM, if one considers that the key to super-resolution is the control and acquisition of some on/off behavior of the fluorescent labels [51], then it appears that event-based sensors are particularly well suited for the purpose of blinking-based imaging of continuous structures since it essentially detects only the on and off transitions. In the same situation, frame-based approaches usually run extra steps aiming at extracting dynamic components such as median temporal filtering or molecule linking over several frames, adding complexity and noise or errors. Furthermore, these steps usually fail under challenging imaging conditions such as when the density of PSFs is high. On the contrary, event-based SMLM is largely unaffected in these conditions, which we attribute to both its ability to extract the useful information only, and the possibility to easily perform efficient temporal resampling, reducing signal density to manageable levels.

We expect that event-based SMLM could benefit from several improvements, both from the point of view of the optical design and that of the processing software. Strategies could be implemented to adapt the spatio-temporal analysis (i.e. the area of the PSF analysis and the processing time base) to the local density (in space and time) of the output signal. This would probably lead to an improvement in calculation speed and localization performance. Improvements could also be made using better suited PSF models for the position calculation [52, 53, 54, 42].

Given the high density imaging capabilities, we also hypothesize that event-based SMLM could be used for quantitative applications in dSTORM since the molecules blinking during the pumping phase can be effectively counted. In other cases where acquisition speed matters most (such as for live imaging or for high content screening), it could also benefit dSTORM and PALM experiments by allowing faster acquisitions, as super-resolved images can be obtained in a few minutes only.

Another very different set of applications where event-based SMLM may excel could be modulated excitation techniques. This can give access to a parameter of interest such as the excitation wavelength in simultaneous multicolor experiments [55], the lateral [56, 57] or axial [5, 58] positions, or the orientation of the dipole associated with the molecule [59]. In all these cases, however, the temporal aspect is crucial. While some approaches apply very low exposure times at the cost of low signal, others use pre-demodulated detection paths in combination with fast switching between the channels, which brings experimental complexity and reduces the effective field of view. We hypothesize that given their speed event-based sensors should be readily able to record the temporal sampling-free dynamics of the modulated response without reducing the signal level by splitting the useful photons.

## Acknowledgements

We acknowledge Christian Hubert as well as Tual Monfort and Olivier Thouvenin for their invaluable help with conceiving the project, as well as for hardware loan. We also thank R. Margoth Córdova-Castro, Vittore Scolari, Antoine Coulon, Sandrine Lévêque-Fort and Emmanuel Fort for fruitful discussions and feedback.

## Author contributions

C.C. and I.I. conceived the project and C.C. led its development, supervised by I.I. C.C. designed the optical setup and processing algorithms, and performed the acquisitions, processing and data analysis. C.S. prepared the fixed cells samples and C.C. performed the immunolabeling. C.C. wrote the manuscript original draft, which was reviewed by C.S. and I.I.

## Methods

### Optical setup

A schematic of the optical setup used is presented in **Fig. 1d**. We used a custom-built microscope with a RM21 body and a MANNZ micro- and nano-positioner (Mad City Labs). The illumination and fluorescence collection was done with a Nikon 100x 1.49NA APO TIRF SR oil immersion objective. The excitation was performed thanks to a 638 nm laser (LBX-638-180, 180 mW, Oxxius) and a 488 nm laser (LBX-488-100, 100 mW, Oxxius) with a 405 nm laser for pumping (LBX-405-50, 50 mW, Oxxius). A full multiband filter set (LF405/488/561/635-A-000, Semrock) was used. The excitation consisted of a standard vertical Gaussian beam without any scanning, speckle removal or optical sectioning (except for the DNA-PAINT acquisitions presented in **Fig. 1b–e**, which were performed under TIRF excitation. The fluorescence was sent in the detection module and recorded on the EM-CCD camera (iXon Ultra 897, Andor) and/or on the event-based sensor (EVK V2 Gen4.1, Prophesee). For the DNA-PAINT acquisitions only, the EMCCD as replaced with an sCMOS camera (Prime 95B, Photometrics). Both sensors had their focal planes approximately matched (below 200 nm difference). We used afocal doublets to adjust the pixel sizes to 107 nm (EMCCD/sCMOS) and 67 nm (event-based sensor) in the object plane. Depending on the acquisition, we placed different elements in the detection path thanks to a flip platform—either a 50:50 non-polarizing beamsplitter (Thorlabs), or a a 30(EMCCD):70(event-based sensor) non-polarizing beamsplitter (Thorlabs), or a plane mirror (Thorlabs) to deflect all the signal on the event-based sensor, or nothing to collect all the fluorescence on the EMCCD/sCMOS camera.

### Fluorescent beads sample preparation (Fig. 1g and Fig. 2b–d)

The sample was prepared by diluting dark red (660/680) 40 nm fluorescent beads (F10720, Thermo Fisher) with a dilution factor of 10^-7^ in phosphate buffered saline (PBS) and allowing them to deposit on a coverslip.

The sample used for **Fig. 1e–f** was prepared by diluting dark red (660/680) 200 nm fluorescent beads (F8807, Thermo Fisher) with a dilution factor of 10^-3^ in PBS and allowing them to deposit on a coverslip.

The sample used for **Fig. 4a** was prepared by diluting dark red (660/680) 40 nm fluorescent beads (F10720, Thermo Fisher) with a dilution factor of 8 × 10^-8^ and yellow-green (505/515) 40 nm fluorescent beads (F10720, Thermo Fisher) with a dilution factor of 8 × 10^-6^ in PBS and allowing them to deposit on a coverslip.

### Alexa Fluor 647 on a coverslip sample preparation (Fig. 2e–g)

The sample was prepared by depositing 1.5 μl of the initial solution of AF647 goat anti-mouse antibody (A21237, Thermo Fisher), allowing 5 minutes for the molecules to deposit before rinsing with H_2_O and adding dSTORM buffer. The buffer was composed of 100 mg/ml glucose, 3.86 mg/ml MEA, 0.5 mg/ml glucose oxidase and 1.18 μl/ml catalase in PBS. The sample was illuminated with a 638 nm continuous excitation at an irradiance of 5 kW/cm^2^.

### Biological samples preparation

African green monkey kidney cells (COS-7) were cultured at 37°C and 5 % CO_2_ in DMEM medium containing glutamax (Gibco No. 31966-047), 10 % fetal bovine serum (FBS, Gibco No. A3840401) and 50 U/ml penicillin and 50 μl/ml streptomycin (Gibco No. 15140-148). For experiments, cells were plated on 25 mm diameter glass coverslips (type 1.5) placed in six well plates containing culture medium with 2 % FBS at low density, and fixed on the following day in 0.1 M sodium phosphate buffer (PB), pH 7.4, containing 4 % paraformaldehyde (PFA), 0.2 % glutaraldehyde, 1 % sucrose, at 37°C for 10 minutes, followed by three rinses in PBS. Cells were permeabilized with PBS containing 0.1 % Triton X-100 for 10 minutes and rinsed three times with PBS prior to immunolabeling.

For the dSTORM labeling, the cells were incubated for 1 hour at 37°C with 1:300 mouse anti-*α*-tubulin antibody (Sigma Aldrich, T6199) in PBS + 1 % BSA. This was followed by three washing steps in PBS + 1 % BSA, incubation for 45 minutes at 37°C with 1:300 goat anti-mouse AF647 antibody (Life Technologies, A21237) diluted in PBS + 1 % BSA and three more washes in PBS. Finally, the cells were post-fixed with 3.6 % formaldehyde for 15 min in PBS. The cells were washed in PBS three times and then reduced for 10 minutes with 50 mM NH4Cl (Sigma Aldrich, 254134), followed by three additional washes in PBS.

For the DNA-PAINT labeling, the cells were incubated for 1 hour at 37°C with 1:300 mouse anti-*α*-tubulin antibody (Sigma Aldrich, T6199) in PBS + 1 % BSA. This was followed by three washing steps in PBS + 1 % BSA, incubation for 1 hour at 37°C with 1:200 anti-mouse docking strand 1 antibody (Massive Photonics, MASSIVE-sdAB 2-PLEX) diluted in antibody incubation buffer and three more washes in washing buffer 1x.

### Localization precision and response linearity measurement (Fig. 2b–c)

Acquisitions were performed on fluorescent beads deposited on a coverslip using a square modulated 638 nm excitation with a frequency of 10 Hz and a duty cycle of 0.5. Using a 30:70 (EMCCD:event-based sensor) non-polarizing beamsplitter, acquisitions were captured simultaneously on the camera (100 ms exposure) and on the event-based sensor with various sensitivity values. Event-based frames were generated by summing all the events in Δ*t* =100 ms time bins regardless of their sign. The positions were calculated using a center of mass calculation on both the event-based reconstructed and the camera-acquired frames. Positions were drift-corrected using a direct cross-correlation algorithm. A simple clustering algorithm was used to determine the localization precision from the calculated positions as well as the average numbers of photons or events for each bead. A total of 1000 frames (i.e. 100 s) were included in the statistics. Finally, a registration and colocalization algorithm was used to match the camera-based and the event-based bead positions in order to compare the performances of both methods. The average number of photons per cycle on the camera was calculated for each bead from the camera localization results, and this number was subsequently used to calculate the average number of photons per cycle on the event-based sensor for each bead by applying a ratio corresponding to the beamsplitter ratio (experimentally measured to be 1:2.38).

### Standard density biological experiments (Fig. 3)

EMCCD acquisitions were done with a 30 ms exposure time and a gain of 100, and event-based acquisitions were done with thresholds corresponding to the level of sensitivity called ‘high’ in the characterization. The detection path used either a 50:50 non-polarizing beamsplitter cube (**Fig. 3a–c,f–g**) or a mirror to send all the photons on the event-based sensor (**Fig. 3d–e**). Both sensors were not synchronized but dual-view acquisitions were started and stopped almost simultaneously. We used a dSTORM buffer composed of 100 mg/ml glucose, 3.86 mg/ml MEA, 0.5 mg/ml glucose oxidase and 1.18 μl/ml catalase in PBS and a 638 nm continuous excitation with an irradiance of 5 kW/cm^2^. After a pumping phase of a few minutes, the acquisitions were started, and stopped after 25 minutes. A low power continuous 405 nm excitation was also added during the second half of the acquisition to increase the density of detections.

EMCCD frames were processed using a wavelet algorithm (similar to [60]) to detect the PSFs and each PSF was localized on a ±4 pixels area centered around the maximum. Positions were estimated using either Gaussian fitting or center of mass calculation. Drift was corrected using a direct cross-correlation algorithm using the sample itself as a reference (no fiducial markers were used).

Event-based data were processed as follows. Positive events were binned in Δ*t*=20 ms frames, on which PSFs were detected using a wavelet algorithm. This yielded the rough reference space and time positions *x*_0_, *y*_0_ and *T_0_* for each PSF. Molecules were then localized from a subset of all the events (both positive and negative) corresponding to an area of ±4 pixels around (*x*_0_, *y*_0_) and to times in the interval [*T*_0_ – 60 ms, *T*_0_ + 120 ms]. The position of the center was estimated using a center of mass calculation. Other data were extracted—the total number of events N in the subset, the time of the rising edge *t*_+_ (taken as the mean time of the positive events) and the time of the falling edge *t*_−_ (taken as the mean time of the negative events). These were used to calculate the ON time of each molecule *t_ON_* = *t*_−_ – *t*_+_. Positions were drift corrected using the same direct cross-correlation as for the frame-based data.

### High density biological experiments (Fig. 4b–h)

High density DNA-PAINT acquisitions (**Fig. 4b–e**) were done with an sCMOS camera set to 50 ms exposure time, and the event-based sensor with a sensitivity corresponding to the level called ‘high’ in the characterization. All the photons were sent sequentially to the sCMOS camera first, then to the event-based sensor, taking advantage of the resistance of DNA-PAINT to photobleaching. We used an imager 1 concentration of 8 nM and a 638 nm continuous excitation with an irradiance of 0.5 kW/cm^2^ in TIRF configuration. A pumping phase was allowed to stabilize the density of PSFs before starting the acquisition, and data was recorded during 10 minutes for each sensor.

High density dSTORM acquisitions (**Fig. 4f–h**) were done with an EMCCD camera set to 30 ms exposure time and a gain of 100, and with the event-based sensitivity corresponding to ‘high’ in the characterization. The detection path used a 50:50 non-polarizing beamsplitter cube for simultaneous EMCCD/event-based acquisition. Both sensors were not synchronized but dual-view acquisitions were started and stopped almost simultaneously. We used a dSTORM buffer composed of 100 mg/ml glucose, 3.86 mg/ml MEA, 0.5 mg/ml glucose oxidase and 1.18 μl/ml catalase in PBS and a 638 nm continuous excitation with an irradiance of 5 kW/cm^2^. No pumping phase was allowed, so the acquisitions were started immediately and stopped after 250 seconds. A low power continuous 405 nm excitation was also used during the entire acquisition to increase the detection density.

In each case, camera frames were processed using a wavelet algorithm to detect the PSFs and each PSF was localized on a ±3 pixels area centered around the maximum. Positions were estimated using either Gaussian fitting or center of mass calculation. Drift was corrected using a direct cross-correlation algorithm using the sample itself as a reference (no fiducial markers were used). Note that, while all the results presented were obtained with our usual custom-written Python processing algorithm, we also tried the widely used ThunderSTORM ImageJ plugin [61], which yielded very similar results (not displayed here).

Event-based data was processed as follows. The dataset was split in two subsets containing the positive and negative events respectively. Positive events were binned in Δ*t*=10 ms frames, on which PSFs were detected using a wavelet algorithm. This yielded the rough reference space and time positions *x*_0_, *y*_0_ and *T_0_* for each PSF. Molecules were then localized from a subset of all the events (both positive and negative) corresponding to an area of ±3 pixels around (*x*_0_, *y*_0_) and to times in the interval [*T*_0_ – 30 ms, *T*_0_ + 30 ms]. The position of the center was estimated using a center of mass calculation or a Gaussian fitting. The total number of events N in the subset corresponding to the molecule was also extracted. The same processing was run on the negative events subset with the same parameters. Due to the separate processing of the positive and negative events, the ON times could not be extracted since we did not use a dedicated program to link the rising and falling edges. Positions were drift corrected using the same direct cross-correlation as for the frame-based data.

### FRC resolution calculation

All FRC resolution measurements were done using the Fiji plugin NanoJ-SQUIRREL [36]. Localization lists were randomly split in two statistically independent sets and super-resolution images were generated with a pixel size of 5 nm. FRC maps were calculated with a number of blocks per axis of 30 (for full images) or 10 (for zooms on small sub-regions). We noticed little variation of the results with the calculation parameters.

### Cramér-Rao Lower Bound calculation (Fig. 2b)

CRLB values were estimated for the EMCCD camera using equation 4 from [62]. The values of the parameters were measured experimentally.

### Localization and data treatment software

The localization and data treatment software is described in **Supplementary note 2.** It should be noted that no filtering was used in the event-based processing. All the processing and rendering was performed using a home-written Python code. The function used to read the event files was borrowed from the Metavision SDK open samples.

## Code availability

Processing codes are made available on Github at the following address: https://github.com/Clement-Cabriel/Evb-SMLM.git

The repository contains the codes for reading files and converting them into frame stacks, and the single-molecule localization code will be added soon. More codes will be added in the future. Datasets are available on the same repository to test the codes.

## Data availability

Dataset examples are available on the Github repository (see **Code availability**). More datasets are available from the corresponding author upon reasonable request.

## Supplementary material

**Supplementary note 1:** Detailed working principle of the sensor’s response.

**Supplementary note 2:** Event-based single-molecule processing algorithms.

**Supplementary Note 3:** Event-based temporal resampling of the signal and PSF density.

**Supplementary Figure 1:** FRC resolution measurements on the acquisitions presented in **Fig. 3** (standard density dSTORM).

**Supplementary Figure 2:** Frames extracted from the acquisition presented in **Fig. 4** (high density dSTORM).

**Supplementary Table 1:** FRC resolution measurements on all the biological samples presented.

### Supplementary note 1: Detailed working principle of the sensor’s response

An event-based sensors is a rectangular or square array of pixels that are sensitive to optical intensity variations only. Since each pixel works independently of the rest of the array, the principle of the sensor can be described through the response of one pixel.

At each time *t*, the light intensity *I*(*t*) (i.e. number of photons incoming on the pixel per unit time) is linearly converted into an electronic response. The pixel stores a reference level *I*_ref_(*t*), which is then used to detect meaningful variations of the intensity relative to this reference level. Two thresholds *B*_+_ and *B*_−_ are used to set the sensitivity and can be modified. They are used respectively to detect an increase and decrease in intensity. While *B*_+_ and *B*_−_ can be set to different values (all pixels use the same values of *B*_+_ and *B*_−_, which are set as input parameters), we always use symmetric values (i.e. *B*_−_ = 1/*B*_+_) in this work. According to the specifications given by the manufacturer, the intensity variation detection is logarithmic rather than linear. Therefore, a signal is triggered as soon as *I*(*t*)/*I*_ref_(*t*) > *B*_+_ (positive event) or *I*(*t*)/*I*_ref_(*t*) < *B*_−_ (negative event). The principle of the response of a pixel is illustrated in **Fig. 1a**, and **Fig. 1b** shows how varying the values of the sensitivity *B* changes the output signal. From a quantitative point of view, these thresholds can be as low as 11 % of the reference level [32].

Whenever an event is triggered, the sensor returns a signal containing the *x* and *y* pixel coordinates, the polarity (positive or negative) and the timestamp of the trigger. These coordinates are added to the output file. Note that the output is essentially a sparse list rather than a dense matrix as in the case of a frame-based camera.

After the event is triggered, the value of the reference level is replaced with the value of the intensity at the time of the trigger. As a consequence, a large variation of the signal causes the pixel to return several events in a quick succession, which readily provides a certain *quantitativity* in the detection. Finally, it should be noted that the sensor has a dead time in the range of a few μs to 100 μs (which is larger for lower intensities), which effectively sets an upper limit to its bandwidth. During this dead time, the pixel cannot generate more events but the reference level is stored. This means that for a large variation of intensity, several events are returned, separated in time by the dead time value, even if the variation is instantaneous.

### Supplementary note 2: Event-based single-molecule processing algorithms

Our processing approach to single-molecule localization is partly frame-based and partly event list-based. More precisely, frames are generated to detect the spots, and the localization is then performed only on the event list.

#### Standard density localization

For standard density acquisitions, events from the raw output list are binned into frames with a given Δ*t* (typically between 10 ms and 50 ms), which are used for PSF detection. This step is done on the positive events only (i.e. when the molecule turns on) using a wavelet decomposition algorithm described in [60]. Then, each molecule is localized using a subset of events corresponding to a spatial ROI of ±4 pixels (i.e. a total area of 9×9 pixels = 600×600 nm) around the center of the PSF, and a temporal ROI of [*T*_0_ – *δT*_start_, *T*_0_ + *δT*_end_ around the timestamp of the frame *T*_0_, with the (negative) lower boundary *δT*_start_ around 20 ms to 60 ms, and the upper boundary *δT*_end_ around 80 ms to 200 ms (*δT*_end_ has to be sufficient to cover the full ON time of the molecule). In this subset of events, the *x* and *y* positions are determined by calculating the center of mass of all the events or by performing a Gaussian fitting of the PSF. Moreover, complementary information can also be extracted, such as the number of positive and negative events *N*_+_ and *N*_−_, the time when the molecule turns on *t*_+_ (taken as the mean of the times of the positive events) and the time when it turns off (mean of the times of the negative events) and the ON time, defined as *t_ON_* = *t*_−_ – *t*_+_. Finally, the drift is corrected using a direct cross correlation algorithm (in other words, the reference is the sample image itself rather than fiducial markers).

#### High density localization

The event-based processing code is adapted to resample the data on shorter time scales for high density imaging (dSTORM or DNA-PAINT). More specifically, frames are generated from the positive events only with a time bin of Δ*t*=10 ms, and the localization is subsequently performed on the corresponding positive-only events within a short time range [*T*_0_ – 30 ms, *T*_0_ + 30 ms] centered around the frame timestamp *T*_0_. By running the processing on such a fine time scale, we extract only the moment when a molecule turns on while its neighbours maintain a roughly constant level of emission. Then, a similar processing is performed on the ngative events to extract the moment when the molecules turn off. Even though this loses track of the link between the rising edge and the falling edge for each molecule and thus prevents the extraction of the ON time, it retains the quantitative aspect of the measurement (each rising edge generates one positive localization, and each falling edge generates one negative localization), so molecules can still be counted. The rest of the localization workflow is unchanged compared to the standard density processing.

### Supplementary note 3: Event-based temporal resampling of the signal and PSF density

In the context of frame-based SMLM, the density of PSFs per frame is determined by several factors. The density of the protein imaged and the density of the fluorescent labelling can be considered together as the first. The second is the temporal dynamics of the blinking. The higher the ratio between the ON and the OFF times of a molecule, the higher the density. The last factor is the exposure time—indeed, if it is larger than the ON time of the molecules, then the density of PSFs per frame increases. However, the exposure time is usually set to a value close to or slightly lower than the ON time of the molecules (in particular, this was the case in this work). Therefore, the density of PSFs is entirely determined by the former two parameters, i.e. density of the fluorescent molecules and the blinking dynamics. To reduce the density, it is possible to decrease the photoactivation rate in PALM, or the concentration of imager strands in DNA-PAINT. This comes at the cost of longer acquisitions, however, which can be a problem in living samples or high throughput imaging. In dSTORM, there is no such straightforward way to adapt the blinking dynamics.

With an event-based sensor, on the contrary, the ON time does not matter much. Indeed, if one molecule turns on, then it does not need to turn off before another can be localized in the same diffraction-limited volume. Rather than the ON time, the density of PSFs is determined by the time it takes for most of the events in a PSF to be returned. As illustrated in **Fig. 1f** and **Fig. 2g**, it is around 10–30 ms, and we suspect that it could be reduced by carefully tuning the electronic parameters of the sensor.

In comparison, for the DNA-PAINT results displayed in **Fig. 4b–e**, each imager strand stays bound for several hundreds of ms on average. The event-based sensing readily provides a reduction of the spatio-temporal density of signal by a factor 10, therefore allowing standard density processing approaches to perform well.

Similarly, dSTORM acquisitions (**Fig. 4f–h**) can benefit greatly from the event-based approach for high density acquisitions. **Supplementary Fig. 2** shows a ROI of an EMCCD frame, where the PSFs overlap significantly, and an event-based generated frame of the positive only events in a Δ*t*=10 ms time bin. On that frame (used only for the PSF detection, see **Methods**), the density of molecules is vastly reduced, which means that although many neighbouring molecules are in an ON state, the variations of the signal are sparse enough for the PSFs to be spotted individually. Note that the EMCCD exposure time (30 ms) is below the typical ON times of the dyes, which implies that the PSF cannot be made sparser by increasing the framerate, as the density is mostly determined by the density of the target protein, the density of labeling as well as the global fraction of active dyes (in other words the pumping rate and elapsed pumping time).

It should be noted that our approach fundamentally does not rely on high density algorithms since the event-based sensing readily provides sparse PSFs in cases where the sparsity criterion is not achieved with scientific cameras.

**Supplementary Figure 1:**
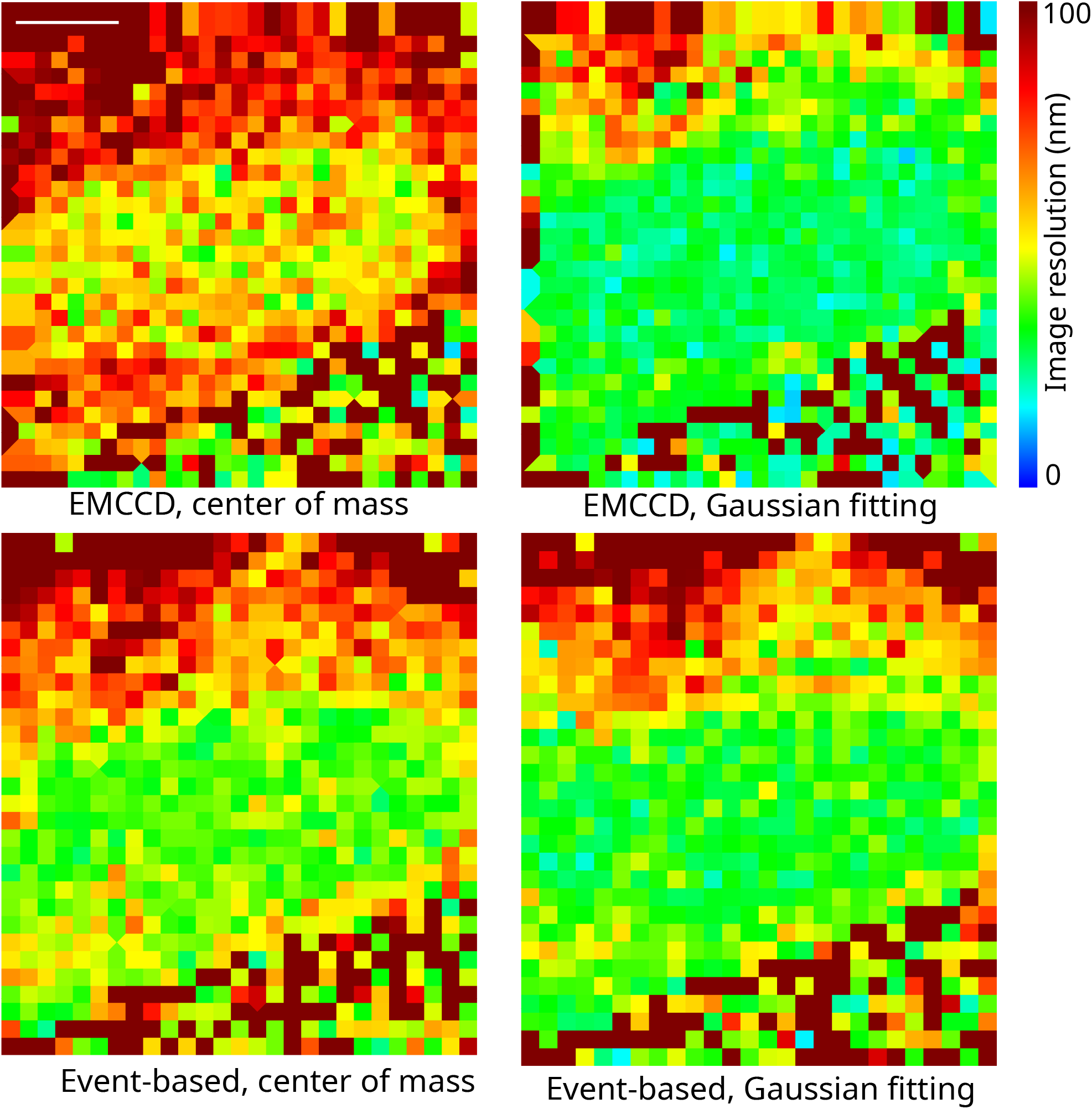
FRC resolution measurements on fixed COS-7 cells labeled with AF647 against *α*-tubulin imaged in standard density dSTORM with the event-based sensor and the EM-CCD camera using a 50:50 beamsplitter in the detection path (corresponding to the data displayed in **Fig. 3)**. Resolution maps were obtained using the NanoJ-SQUIRREL Fiji plugin [36] (see **Methods**). Scale bar: 5 μm.

**Supplementary Figure 2:**
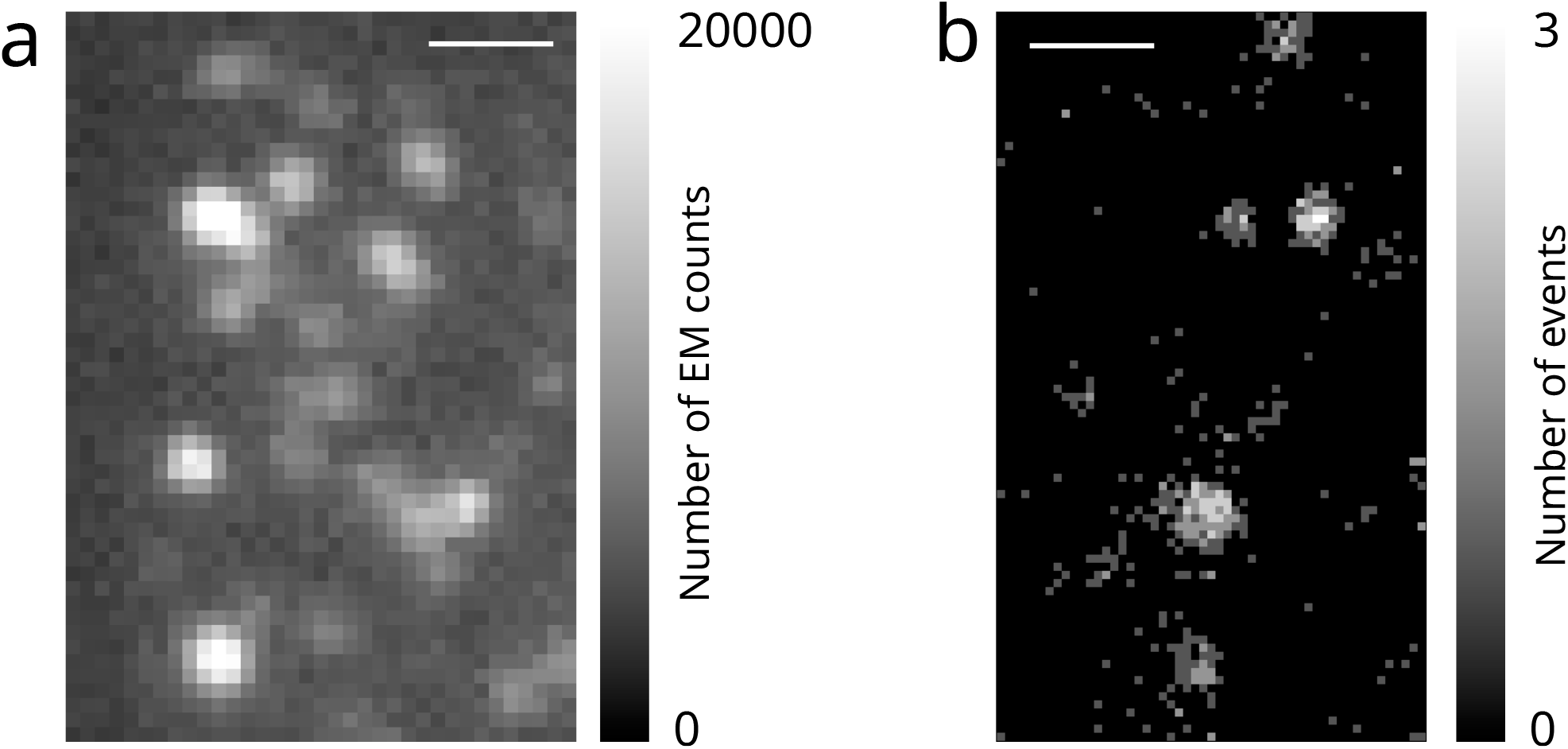
Frames extracted from an acquisition on fixed COS-7 cells labeled with AF647 against *α*-tubulin acquired with a 50:50 beamsplitter over 250 seconds in a dense regime where the PSF exhibit noticeable overlap (presented in **Fig. 4f–h**). **a** Single 30 ms exposure frame taken from the EMCCD blinking movie. **b** Single frame (corresponding to a slightly different instant in the acquisition as **a**) generated from all the positive events detected in Δ*t* = 10 ms (this frame is used only for the PSF detection). Scale bars: 1 μm.

**Supplementary Table 1:** FRC resolution measurements on fixed COS-7 cells labeled with AF647 against *α*-tubulin under various acquisition conditions.

| Acquisition conditions | Corresponding figure | Sensor and localization method | Average resolution (nm) |
| --- | --- | --- | --- |
| Standard density, full field, 50:50 beamsplitter | **Fig. 3a** | Event-based, center of mass<br>Event-based, Gaussian fitting<br>EMCCD, center of mass<br>EMCCD, Gaussian fitting | 36<br>30<br>50<br>28 |
| Standard density, zoom, 50:50 beamsplitter | **Fig. 3b** | Event-based, center of mass<br>Event-based, Gaussian fitting<br>EMCCD, center of mass<br>EMCCD, Gaussian fitting | 34<br>28<br>27<br>44 |
| Standard density, full field, 100 % on the event-based sensor | **Fig. 3d left** | Event-based, center of mass<br>Event-based, Gaussian fitting | 28<br>24 |
| Standard density, zoom, 100 % on the event-based sensor | **Fig. 3d right** | Event-based, center of mass<br>Event-based, Gaussian fitting | 28<br>23 |
| High density dSTORM, full field, 50:50 beamsplitter | **Fig. 4f,h** | Event-based, center of mass<br>Event-based, Gaussian fitting<br>EMCCD, center of mass<br>EMCCD, Gaussian fitting | 60<br>58<br>136<br>92 |
| High density dSTORM, zoom, 50:50 beamsplitter | **Fig. 4g** | Event-based, center of mass<br>Event-based, Gaussian fitting<br>EMCCD, center of mass<br>EMCCD, Gaussian fitting | 52<br>50<br>114<br>84 |

## References

[1] E. Betzig, G. H. Patterson, R. Sougrat, O. W. Lindwasser, S. Olenych, J. S. Bonifacino, M. W. Davidson, J. Lippincott-Schwartz, and H. F. Hess, “Imaging intracellular fluorescent proteins at nanometer resolution.,” Science (New York, N.Y.), vol. 313, pp. 1642–5, sep 2006.

[2] S. T. Hess, T. P. K. Girirajan, and M. D. Mason, “Ultra-high resolution imaging by fluorescence photoactivation localization microscopy.,” Biophysical journal, vol. 91, pp. 4258–72, dec 2006.

[3] M. J. Rust, M. Bates, and X. Zhuang, “Sub-diffraction-limit imaging by stochastic optical reconstruction microscopy (STORM),” Nature Methods, vol. 3, pp. 793–796, aug 2006.

[4] F. Balzarotti, Y. Eilers, K. C. Gwosch, A. H. Gynnå, V. Westphal, F. D. Stefani, J. Elf, and S. W. Hell, “Nanometer resolution imaging and tracking of fluorescent molecules with minimal photon fluxes,” Science, vol. 355, pp. 606–612, dec 2016.

[5] P. Jouchet, C. Cabriel, N. Bourg, M. Bardou, C. Poüs, E. Fort, and S. Lévêque-Fort, “Nanometric axial localization of single fluorescent molecules with modulated excitation,” Nature Photonics, vol. 15, pp. 297–304, jan 2021.

[6] G. V. Los, L. P. Encell, M. G. McDougall, D. D. Hartzell, N. Karassina, C. Zimprich, M. G. Wood, R. Learish, R. F. Ohana, M. Urh, D. Simpson, J. Mendez, K. Zimmerman, P. Otto, G. Vidugiris, J. Zhu, A. Darzins, D. H. Klaubert, R. F. Bulleit, and K. V. Wood, “HaloTag: A novel protein labeling technology for cell imaging and protein analysis,” ACS Chemical Biology, vol. 3, no. 6, pp. 373–382, 2008.

[7] P. J. Bosch, I. R. Corrêa, M. H. Sonntag, J. Ibach, L. Brunsveld, J. S. Kanger, and V. Subramaniam, “Evaluation of fluorophores to label SNAP-tag fused proteins for multicolor single-molecule tracking microscopy in live cells.,” Biophysical journal, vol. 107, pp. 803–14, aug 2014.

[8] M. Mikhaylova, B. M. C. Cloin, K. Finan, R. van den Berg, J. Teeuw, M. M. Kijanka, M. Sokolowski, E. a. Katrukha, M. Maidorn, F. Opazo, S. Moutel, M. Vantard, F. Perez, P. M. P. van Bergen en Henegouwen, C. C. Hoogenraad, H. Ewers, and L. C. Kapitein, “Resolving bundled microtubules using anti-tubulin nanobodies,” Nature Communications, vol. 6, no. May, p. 7933, 2015.

[9] L. D. Lavis, “Teaching old dyes new tricks: Biological probes built from fluoresceins and rhodamines,” Annual Review of Biochemistry, vol. 86, pp. 825–843, jun 2017.

[10] J. B. Grimm, T. A. Brown, B. P. English, T. Lionnet, and L. D. Lavis, “Synthesis of janelia fluor HaloTag and SNAP-tag ligands and their use in cellular imaging experiments,” in Methods in Molecular Biology, pp. 179–188, Springer New York, 2017.

[11] M. Lelek, M. T. Gyparaki, G. Beliu, F. Schueder, J. Griffié, S. Manley, R. Jungmann, M. Sauer, M. Lakadamyali, and C. Zimmer, “Single-molecule localization microscopy,” Nature Reviews Methods Primers, vol. 1, jun 2021.

[12] Y.-L. Wu, A. Tschanz, L. Krupnik, and J. Ries, “Quantitative data analysis in single-molecule localization microscopy,” Trends in Cell Biology, vol. 30, pp. 837–851, nov 2020.

[13] B. Huang, W. Wang, M. Bates, and X. Zhuang, “Three-Dimensional Super-Resolution Imaging by Stochastic Optical Reconstruction Microscopy,” Science, vol. 319, no. 5864, pp. 810–813, 2008.

[14] S. R. P. Pavani, M. A. Thompson, J. S. Biteen, S. J. Lord, N. Liu, R. J. Twieg, R. Piestun, and W. E. Moerner, “Three-dimensional, single-molecule fluorescence imaging beyond the diffraction limit by using a double-helix point spread function,” Proceedings of the National Academy of Sciences, vol. 106, pp. 2995–2999, feb 2009.

[15] N. Bourg, C. Mayet, G. Dupuis, T. Barroca, P. Bon, S. Lécart, E. Fort, and S. Lévêque-Fort, “Direct optical nanoscopy with axially localized detection,” Nature Photonics, no. August, 2015.

[16] C. Cabriel, N. Bourg, P. Jouchet, G. Dupuis, C. Leterrier, A. Baron, M.-A. Badet-Denisot, B. Vauzeilles, E. Fort, and S. Lévêque-Fort, “Combining 3d single molecule localization strategies for reproducible bioimaging,” Nature Communications, vol. 10, apr 2019.

[17] S. A. Maynard, P. Rostaing, N. Schaefer, O. Gemin, A. Candat, A. Dumoulin, C. Villmann, A. Triller, and C. G. Specht, “Identification of a stereotypic molecular arrangement of endogenous glycine receptors at spinal cord synapses,” eLife, vol. 10, dec 2021.

[18] I. M. Khater, I. R. Nabi, and G. Hamarneh, “A review of super-resolution single-molecule localization microscopy cluster analysis and quantification methods,” Patterns, vol. 1, p. 100038, jun 2020.

[19] H. Verdier, F. Laurent, A. Cassé, C. L. Vestergaard, C. G. Specht, and J.-B. Masson, “A maximum mean discrepancy approach reveals subtle changes in α-synuclein dynamics,” bioRxiv, apr 2022.

[20] A. Lampe, V. Haucke, S. J. Sigrist, M. Heilemann, and J. Schmoranzer, “Multi-colour direct STORM with red emitting carbocyanines,” Biology of the Cell, vol. 104, no. 4, pp. 229–237, 2012.

[21] Z. Zhang, S. J. Kenny, M. Hauser, W. Li, and K. Xu, “Ultrahigh-throughput single-molecule spectroscopy and spectrally resolved super-resolution microscopy,” Nature Methods, vol. 12, pp. 935–938, aug 2015.

[22] C. A. Valades Cruz, H. A. Shaban, A. Kress, N. Bertaux, S. Monneret, M. Mavrakis, J. Savatier, and S. Brasselet, “Quantitative nanoscale imaging of orientational order in biological filaments by polarized superresolution microscopy,” Proceedings of the National Academy of Sciences, p. 201516811, 2016.

[23] A. Kinkhabwala, Z. Yu, S. Fan, Y. Avlasevich, K. Müllen, and W. E. Moerner, “Large single-molecule fluorescence enhancements produced by a bowtie nanoantenna,” Nature Photonics, vol. 3, pp. 654–657, oct 2009.

[24] E. Lerner, T. Cordes, A. Ingargiola, Y. Alhadid, S. Chung, X. Michalet, and S. Weiss, “Toward dynamic structural biology: Two decades of single-molecule förster resonance energy transfer,” Science, vol. 359, jan 2018.

[25] D. Bouchet, J. Scholler, G. Blanquer, Y. D. Wilde, I. Izeddin, and V. Krachmalnicoff, “Probing near-field light–matter interactions with single-molecule lifetime imaging,” Optica, vol. 6, p. 135, jan 2019.

[26] G. Blanquer, B. van Dam, A. Gulinatti, G. Acconcia, Y. D. Wilde, I. Izeddin, and V. Krachmalnicoff, “Relocating single molecules in super-resolved fluorescence lifetime images near a plasmonic nanostructure,” ACS Photonics, vol. 7, pp. 393–400, jan 2020.

[27] A. F. Koenderink, R. Tsukanov, J. Enderlein, I. Izeddin, and V. Krachmalnicoff, “Super-resolution imaging: when biophysics meets nanophotonics,” Nanophotonics, vol. 11, pp. 169–202, dec 2021.

[28] N. Oleksiievets, Y. Sargsyan, J. C. Thiele, N. Mougios, S. Sograte-Idrissi, O. Nevskyi, I. Gregor, F. Opazo, S. Thoms, J. Enderlein, and R. Tsukanov, “Fluorescence lifetime DNA-PAINT for multiplexed super-resolution imaging of cells,” Communications Biology, vol. 5, jan 2022.

[29] A. Beghin, A. Kechkar, C. Butler, F. Levet, M. Cabillic, O. Rossier, G. Giannone, R. Galland, D. Choquet, and J.-B. Sibarita, “Localization-based super-resolution imaging meets high-content screening,” Nature Methods, vol. 14, pp. 1184–1190, oct 2017.

[30] A. E. S. Barentine, Y. Lin, E. M. Courvan, P. Kidd, M. Liu, L. Balduf, T. Phan, F. Rivera-Molina, M. R. Grace, Z. Marin, M. Lessard, J. R. Chen, S. Wang, K. M. Neugebauer, J. Bewersdorf, and D. Baddeley, “An integrated platform for high-throughput nanoscopy,” bioRxiv, apr 2019.

[31] F. Liao, F. Zhou, and Y. Chai, “Neuromorphic vision sensors: Principle, progress and perspectives,” Journal of Semiconductors, vol. 42, p. 013105, jan 2021.

[32] G. Gallego, T. Delbruck, G. Orchard, C. Bartolozzi, B. Taba, A. Censi, S. Leutenegger, A. J. Davison, J. Conradt, K. Daniilidis, and D. Scaramuzza, “Event-based vision: A survey,” IEEE Transactions on Pattern Analysis and Machine Intelligence, vol. 44, pp. 154–180, jan 2022.

[33] “Prophesee gen4.1 product brief: https://support.prophesee.ai/portal/en/kb/articles/evk4-hd-product-brief.”

[34] D. A. Helmerich, G. Beliu, D. Taban, M. Meub, M. Streit, A. Kuhlemann, S. Doose, and M. Sauer, “Photoswitching fingerprint analysis bypasses the 10-nm resolution barrier,” Nature Methods, vol. 19, pp. 986–994, aug 2022.

[35] R. P. J. Nieuwenhuizen, K. A. Lidke, M. Bates, D. L. Puig, D. Grünwald, S. Stallinga, and B. Rieger, “Measuring image resolution in optical nanoscopy,” Nature Methods, vol. 10, pp. 557–562, apr 2013.

[36] S. Culley, D. Albrecht, C. Jacobs, P. M. Pereira, C. Leterrier, J. Mercer, and R. Henriques, “Quantitative mapping and minimization of super-resolution optical imaging artifacts,” Nature Methods, vol. 15, pp. 263–266, feb 2018.

[37] O. K. Wade, J. B. Woehrstein, P. C. Nickels, S. Strauss, F. Stehr, J. Stein, F. Schueder, M. T. Strauss, M. Ganji, J. Schnitzbauer, H. Grabmayr, P. Yin, P. Schwille, and R. Jungmann, “124-color super-resolution imaging by engineering DNA-PAINT blinking kinetics,” Nano Letters, vol. 19, pp. 2641–2646, mar 2019.

[38] J. E. Donehue, E. Wertz, C. N. Talicska, and J. S. Biteen, “Plasmon-enhanced brightness and photostability from single fluorescent proteins coupled to gold nanorods,” The Journal of Physical Chemistry C, vol. 118, pp. 15027–15035, jun 2014.

[39] T. Dertinger, R. Colyer, G. Iyer, S. Weiss, and J. Enderlein, “Fast, background-free, 3d super-resolution optical fluctuation imaging (SOFI),” Proceedings of the National Academy of Sciences, vol. 106, pp. 22287–22292, dec 2009.

[40] S. Geissbuehler, C. Dellagiacoma, and T. Lasser, “Comparison between SOFI and STORM,” Biomedical Optics Express, vol. 2, p. 408, jan 2011.

[41] S. Hugelier, J. J. de Rooi, R. Bernex, S. Duwé, O. Devos, M. Sliwa, P. Dedecker, P. H. C. Eilers, and C. Ruckebusch, “Sparse deconvolution of high-density super-resolution images,” Scientific Reports, vol. 6, feb 2016.

[42] A. Speiser, L.-R. Müller, P. Hoess, U. Matti, C. J. Obara, W. R. Legant, A. Kreshuk, J. H. Macke, J. Ries, and S. C. Turaga, “Deep learning enables fast and dense single-molecule localization with high accuracy,” Nature Methods, vol. 18, pp. 1082–1090, sep 2021.

[43] R. Wombacher, M. Heidbreder, S. van de Linde, M. P. Sheetz, M. Heilemann, V. W. Cornish, and M. Sauer, “Live-cell super-resolution imaging with trimethoprim conjugates,” Nature Methods, vol. 7, pp. 717–719, aug 2010.

[44] X. Xu, Ke;Babcock, Hazen P; Zhuang, “Dual-objective STORM reveals three-dimensional filament organization in the actin cytoskeleton,” Nature methods, vol. 9, no. 2, pp. 185–188, 2012.

[45] R. Boudjemaa, C. Cabriel, F. Dubois-Brissonnet, N. Bourg, G. Dupuis, A. Gruss, S. Lévêque-Fort, R. Briandet, M.-P. Fontaine-Aupart, and K. Steenkeste, “Impact of bacterial membrane fatty acid composition on the failure of daptomycin to kill staphylococcus aureus,” Antimicrobial Agents and Chemotherapy, vol. 62, jul 2018.

[46] A. Aristov, B. Lelandais, E. Rensen, and C. Zimmer, “ZOLA-3d allows flexible 3d localization microscopy over an adjustable axial range,” Nature Communications, vol. 9, jun 2018.

[47] D. Mahecic, W. L. Stepp, C. Zhang, J. Griffié, M. Weigert, and S. Manley, “Event-driven acquisition for content-enriched microscopy,” Nature Methods, vol. 19, pp. 1262–1267, sep 2022.

[48] J. Alvelid, M. Damenti, C. Sgattoni, and I. Testa, “Event-triggered STED imaging,” Nature Methods, sep 2022.

[49] E. R. Weeks, “Introduction to the colloidal glass transition,” ACS Macro Letters, vol. 6, pp. 27–34, dec 2016.

[50] C. R. Nugent, K. V. Edmond, H. N. Patel, and E. R. Weeks, “Colloidal glass transition observed in confinement,” Physical Review Letters, vol. 99, p. 025702, jul 2007.

[51] S. W. Hell, “Far-field optical nanoscopy,” Science, vol. 316, pp. 1153–1158, may 2007.

[52] H. P. Babcock and X. Zhuang, “Analyzing single molecule localization microscopy data using cubic splines,” Scientific Reports, vol. 7, apr 2017.

[53] E. Nehme, D. Freedman, R. Gordon, B. Ferdman, L. E. Weiss, O. Alalouf, T. Naor, R. Orange, T. Michaeli, and Y. Shechtman, “DeepSTORM3d: dense 3d localization microscopy and PSF design by deep learning,” Nature Methods, vol. 17, pp. 734–740, jun 2020.

[54] N. Boyd, E. Jonas, H. Babcock, and B. Recht, “Deeploco: Fast 3d localization microscopy using neural networks,” bioRxiv, 2018.

[55] P. A. Gómez-García, E. T. Garbacik, J. J. Otterstrom, M. F. Garcia-Parajo, and M. Lakadamyali, “Excitation-multiplexed multicolor superresolution imaging with fm-STORM and fm-DNA-PAINT,” Proceedings of the National Academy of Sciences, vol. 115, pp. 12991–12996, dec 2018.

[56] L. Gu, Y. Li, S. Zhang, Y. Xue, W. Li, D. Li, T. Xu, and W. Ji, “Molecular resolution imaging by repetitive optical selective exposure,” Nature Methods, vol. 16, pp. 1114–1118, sep 2019.

[57] J. Cnossen, T. Hinsdale, R. Ø. Thorsen, M. Siemons, F. Schueder, R. Jungmann, C. S. Smith, B. Rieger, and S. Stallinga, “Localization microscopy at doubled precision with patterned illumination,” Nature Methods, vol. 17, pp. 59–63, dec 2019.

[58] L. Gu, Y. Li, S. Zhang, M. Zhou, Y. Xue, W. Li, T. Xu, and W. Ji, “Molecular-scale axial localization by repetitive optical selective exposure,” Nature Methods, vol. 18, pp. 369–373, apr 2021.

[59] R. Ø. Thorsen, C. N. Hulleman, B. Rieger, and S. Stallinga, “Photon efficient orientation estimation using polarization modulation in single-molecule localization microscopy,” Biomedical Optics Express, vol. 13, p. 2835, apr 2022.

[60] I. Izeddin, J. Boulanger, V. Racine, C. G. Specht, a. Kechkar, D. Nair, a. Triller, D. Choquet, M. Dahan, and J. B. Sibarita, “Wavelet analysis for single molecule localization microscopy.,” Optics express, vol. 20, pp. 2081–95, jan 2012.

[61] M. Ovesný, P. Křížek, J. Borkovec, Z. Švindrych, and G. M. Hagen, “ThunderSTORM: a comprehensive ImageJ plug-in for PALM and STORM data analysis and super-resolution imaging,” Bioinformatics, vol. 30, pp. 2389–2390, may 2014.

[62] S. Stallinga and B. Rieger, “The effect of background on localization uncertainty in single emitter imaging,” in 2012 9th IEEE International Symposium on Biomedical Imaging (ISBI), pp. 988–991, IEEE, may 2012.

